# *In vivo* proteomic mapping through GFP-directed proximity-dependent biotin labelling in zebrafish

**DOI:** 10.1101/2020.11.05.370585

**Authors:** Zherui Xiong, Harriet P. Lo, Kerrie-Ann McMahon, Nick Martel, Alun Jones, Michelle M. Hill, Robert G. Parton, Thomas E. Hall

## Abstract

Protein interaction networks are crucial for complex cellular processes. However, the elucidation of protein interactions occurring within highly specialised cells and tissues is challenging. Here we describe the development, and application, of a new method for proximity-dependent biotin labelling in whole zebrafish. Using a conditionally stabilised GFP-binding nanobody to target a biotin ligase to GFP-labelled proteins of interest, we show tissue-specific proteomic profiling using existing GFP-tagged transgenic zebrafish lines. We demonstrate the applicability of this approach, termed BLITZ (Biotin Labelling In Tagged Zebrafish), in diverse cell types such as neurons and vascular endothelial cells. We applied this methodology to identify interactors of caveolar coat protein, cavins, in skeletal muscle. Using this system, we defined specific interaction networks within *in vivo* muscle cells for the closely related but functionally distinct Cavin4 and Cavin1 proteins.

## Introduction

The understanding of the biological functions of a protein requires detailed knowledge of the molecules with which it interacts. However, robust elucidation of interacting proteins, including not only strong direct protein-protein interactions, but also weak, transient or indirect interactions is challenging. Proximity-dependent biotin labelling (BioID) using genetically engineered biotin ligases has emerged as a novel approach for studying protein-protein interactions and the subcellular proteome in living cells (Roux et al., 2012; Kim et al., 2016; Branon et al., 2018; Ramanathan et al., 2018). When fused to a protein of interest (POI) and expressed in cells, the promiscuous biotin ligases covalently attach biotin to all proteins within a 10 nm radius, which can be subsequently isolated by streptavidin purification and identified by mass spectrometry. Compared with traditional affinity purification with protein-specific antibodies or affinity purification tags, the BioID method has the advantage of being able to capture weak and transient interactions. In addition, unlike conventional methods such as affinity purification, where stringent extraction conditions may disrupt protein-protein interactions, the BioID method does not require proteins to be isolated in their native state. Therefore, harsh protein extraction and stringent wash conditions can be applied, which can improve solubilisation of membrane proteins and reduce false positives (Varnaite and MacNeill, 2016; Gingras et al., 2019).

The BioID method has been widely applied in cell biology to study protein-protein interactions in cultured cells, providing valuable information for building protein interaction networks. However, the reductionist *in vitro* applications described to date, while powerful in their own right, lack the complexity and context to address phenomena that can only be modelled *in vivo*, for example the differentiation of specialised cell types such as those found in muscle, the nervous system and vasculature. The most recent generation of biotin ligases has been applied *in vivo* in invertebrate models; flies (*Drosophila melanogaster*) and worms (*Caenorhabditis elegans*) as well as plants (*Arabidopsis* and *Nicotiana benthamiana*) (Branon et al., 2018; Mair et al., 2019; Zhang et al., 2019). Until now however, the applicability of BioID has been limited by the necessity to genetically tag each POI directly with a biotin ligase and generate transgenic organisms. Here we describe a more versatile approach to the *in vivo* application of BioID in a vertebrate model organism, the zebrafish. Instead of directly fusing the biotin ligase to a POI, we developed a modular system for GFP-directed proteomic mapping by combining BioID with a GFP-binding nanobody (Hamers-Casterman et al., 1993; Rothbauer et al., 2008; Tang et al., 2015; Ariotti et al., 2015). This system couples the power of the BioID system with the ability to use existing GFP-tagged transgenic zebrafish lines for proteomic mapping between different tissues and/or different proteins. We demonstrate the application of this system in screening for proteins associated with the caveolar cast proteins, Cavin1 and Cavin4, in differentiated skeletal muscle which has, to date, been difficult to achieve in culture. These analyses reveal proteins and pathways that are both overlapping and specific to Cavin1 and Cavin4.

## Results

### Proximity biotinylation in live zebrafish embryos

We first tested the ability of a number of biotin ligases to catalyse protein biotinylation in live zebrafish embryos. Initial attempts using BirA* or BioID2 biotin ligases *in vivo* in zebrafish were unsuccessful and resulted in no detectable biotinylation in zebrafish embryos as assessed by streptavidin Western blotting (Supplementary Figure 1). In recent years, the new bioID biotin ligases, BASU, TurboID and miniTurbo, have been developed and showed greatly improved catalytic efficiency and enhanced proximity labelling in cultured cells (Branon et al., 2018; Ramanathan et al., 2018). We therefore tested their ability to catalyse protein biotinylation in live zebrafish embryos. Untagged, cytoplasmically localised biotin ligases were transiently expressed in zebrafish embryos by RNA injection, and the level of protein biotinylation was assessed using streptavidin immunoblot analysis (experimental regimen illustrated in Figure 1A) (Ramanathan et al., 2018; Branon et al., 2018). The biotin ligase mRNA incorporated an EGFP tag for selection of transgene expressing embryos, and a Myc tag for detection of biotin ligase by Western blot. At 24 h post injection, the GFP-positive embryos were dechorionated before incubation in biotin-supplemented media for a further 18 h (Figure 1A). Total protein extracts from fish embryos were then subjected to SDS-PAGE and streptavidin immunoblotting (Figure 1B). TurboID exhibited the strongest biotinylation of endogenous proteins among the three new biotin ligases with 500 μM biotin. Therefore, we chose TurboID for all subsequent experiments. Note the two prominent bands consistently detected around 70 and 135 kDa in all samples likely represent endogenously biotinylated proteins (Housley et al., 2014; Ahmed et al., 2014).

**Figure 1.**
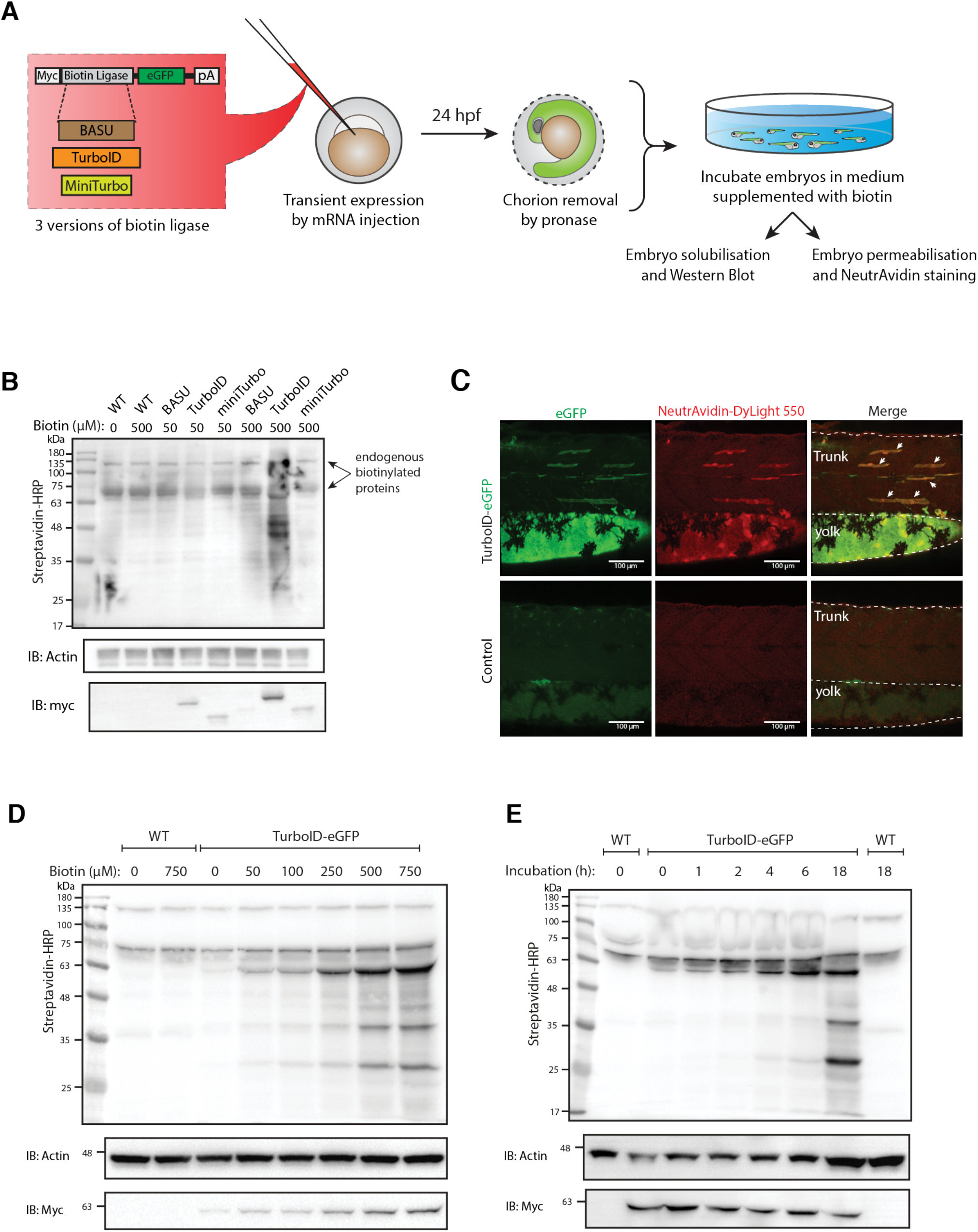
Testing and optimising biotin ligases: BASU, TurboID and miniTurbo. (**A**) A schematic overview of the workflow. The BASU/TurboID/MiniTurbo was transiently expressed in zebrafish embryos by RNA injection. Chorion-removed fish embryos with green fluorescence were selected for incubation in biotin supplemented embryo media for 18 h. After biotin incubation, embryos were analysed by Western blotting and immunofluorescence. (**B**) The streptavidin-HRP blot showing biotinylated proteins in 2 dpf zebrafish embryos expressing eGFP-tagged BASU, TurboID and miniTurbo. Fish embryos were incubated in biotin concentrations of 50 or 500 μM biotin for 18 h before embryo solubilisation and Western blot analysis. Actin immunoblot (IB:Actin) serves as a loading control; the anti-Myc immunoblot (IB:Myc) reflects the protein level of each biotin ligases; each sample is a pool of 30 embryos. (**C**) Representative images of NeutrAvidin staining of biotinylated proteins in 2 dpf zebrafish embryo transiently expressing TurboID-eGFP. Fish muscle and yolk were outlined with dashed lines. White arrows indicate muscle fibres expressing TurboID-eGFP. n=6. Scale bar denotes 100 μm (**D** and **E**) Dependency of TurboID activity on biotin concentration and incubation time. Zebrafish embryos transiently expressing TurboID-eGFP were incubated with embryo media containing 0 to 750 μM biotin for 18 h (**D**) or incubated with 500 μM biotin for 0 to 18 h (**E**) before protein extraction and immunoblot analysis with streptavidin-HRP, anti-Actin and anti-Myc antibodies; each sample is a pool of 30 embryos.

To visualise TurboID-catalysed biotinylation *in situ*, TurboID-expressing embryos were stained with NeutrAvidin-DyLight 550 after biotin incubation (Figure 1C). The mosaic expression of TurboID-GFP in the muscle fibres, as well as expression in the yolk, corresponded with strong NeutrAvidin staining. The mRNA injections frequently gave rise to differing levels of expression between individual muscle cells within the same animal. Therefore, muscle fibres with little or no TurboID-GFP expression served as an internal negative control.

Biotin concentration and incubation time are two crucial factors that affect biotin ligase efficiency in cultured cells (Roux et al., 2012; Kim et al., 2016; Branon et al., 2018; Ramanathan et al., 2018). To achieve the most effective experimental conditions for TurboID application in zebrafish, we sought to optimise these parameters. From our initial experiments with BirA*, we knew that zebrafish embryos are able to tolerate a biotin concentration as high as 800 μM with no obvious morphological abnormalities (Supplementary Figure 2A and B). To determine the optimal biotin concentration for TurboID in zebrafish, TurboID-expressing embryos were incubated in embryo medium containing biotin concentration ranging from 0 to 750 μM for 18 h, followed by lysis and streptavidin immunoblotting. Weak labelling could be seen with 50 μM biotin, increasing through 250 μM, with the strongest labelling at concentration of 500 and 750 μM (Figure 1D). Unlike its application in cultured cells and yeast (Branon et al., 2018), TurboID did not produce detectable exogenous biotinylation without the addition of biotin (Figure 1D). This provides the opportunity for temporal resolution by addition of exogenous biotin at specific developmental stages. Unexpectedly, the anti-Myc immunoblot showed that a higher biotin concentration resulted in more TurboID in the total protein extracts (Figure 1D). Concomitantly, the addition of exogenous biotin did not change the level of endogenous biotinylated proteins in the WT embryos (Figure 1D).

In mammalian cell culture, a 10 min biotin incubation with TurboID is sufficient to visualise biotinylated proteins by immunoblotting and to perform analysis of different organellar proteomes (Branon et al., 2018). However, we did not observe rapid biotinylation in zebrafish within the first 2 h of biotin incubation (Figure 1E). TurboID-induced biotin labelling was only weakly detected after 4 – 6 h incubation and adequate biotinylation was only detected after overnight incubation (18 h).

### *In vivo* proximity biotinylation targeted to a specific subcellular region or a protein of interest

Next, we tested the spatial resolution of TurboID-catalysed biotinylation in zebrafish when TurboID was targeted to a specific subcellular region and to a POI. We tagged TurboID with a nuclear localisation signal (NLS), a plasma membrane localisation motif (CaaX), the transmembrane protein CD44b and the muscle T-tubule enriched membrane protein Cavin4b (Figure 2A). After biotin treatment, the TurboID fusion proteins produced a biotinylation pattern corresponding to the appropriate subcellular location of targeting sequences/proteins in zebrafish embryos (Figure 2B). The spatial resolution of the biotin labelling was remarkable as even the T-tubule structure, which is difficult to resolve in fixed embryos, was clearly visible by NeutrAvidin staining in the embryos expressing Cavin4b-TurboID. Furthermore, the biotinylated protein derived from each TurboID construct gave rise to a unique barcode of protein bands on the streptavidin blot, indicative of proteins specific to each corresponding subcellular compartment (Figure 2C). These results demonstrated that TurboID was able to specifically label a selective subpopulation of endogenous proteins when targeted to a specific subcellular region or protein in zebrafish embryos. Moreover, the TurboID-biotinylated proteins were recoverable from crude fish lysates by affinity purification with streptavidin-conjugated beads (Figure 2D), ready for downstream applications such as identification by mass spectrometry.

**Figure 2.**
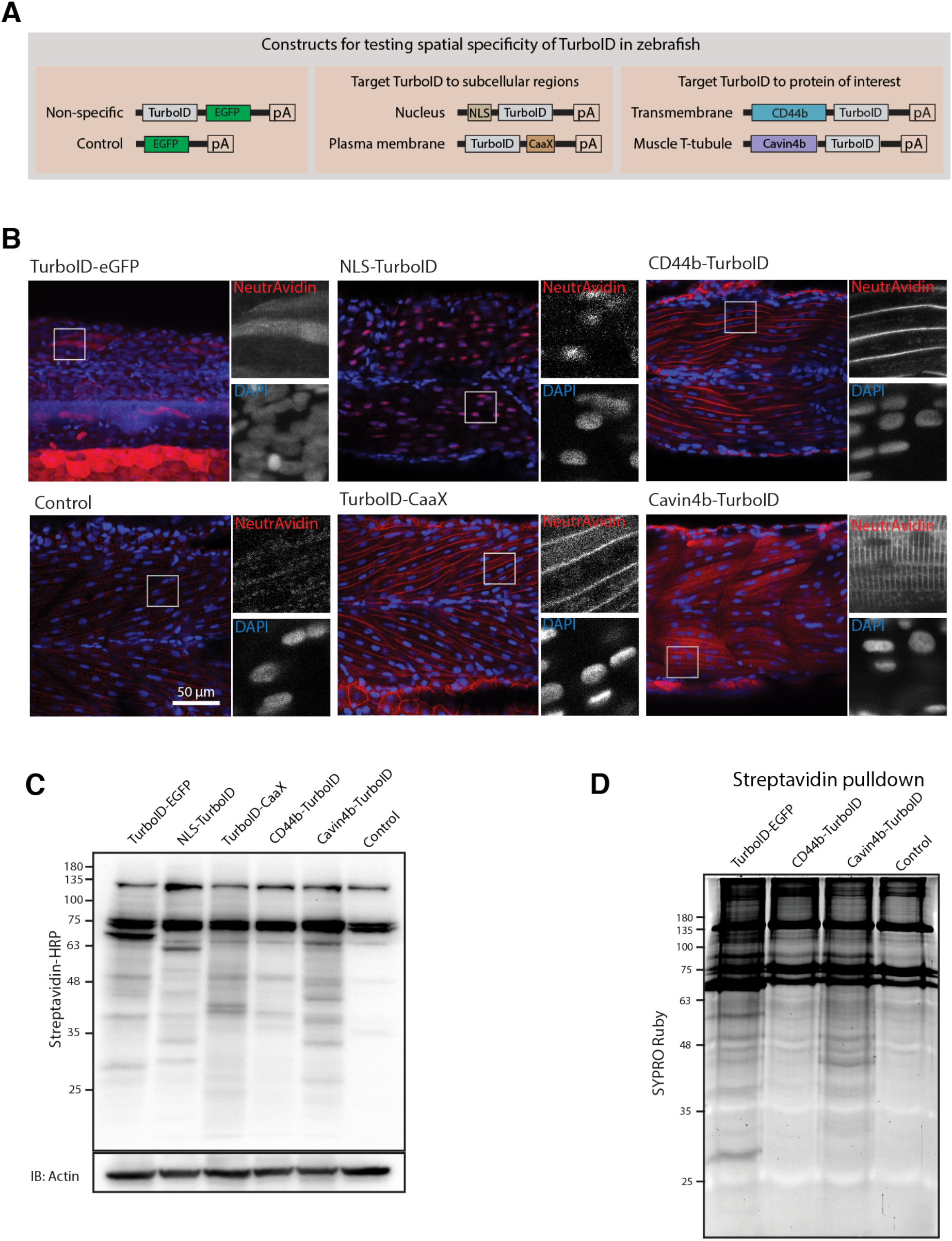
Spatial resolution of TurboID-catalysed biotinylation in zebrafish embryos. (**A**) Schematic representation of eGFP-, NLS-, CaaX-, CD44b- and Cavin4b-tagged TurboID constructs for mRNA injection in zebrafish embryos. TurboID-eGFP was used as a positive control. (**B**) Representative images showing the distribution of biotinylated proteins in 2 dpf zebrafish embryos transiently expressing different TurboID constructs. Negative control fish were injected with eGFP only. Fish embryos were fixed and permeabilised before NeutrAvidin-DyLight staining for biotin and DAPI staining to indicate nuclei. Regions within the white box were magnified and shown in the grey scale for NeutrAvidin and DAPI staining in the right panel; n=3. Scale bar represents 50 μm. (**C**) Streptavidin-HRP blots showing the “protein barcode” produced by biotinylated proteins in fish embryo transiently expressing different TurboID constructs. Actin immunoblot served as a loading control. Each sample is a pool of 30 embryos. (**D**) SYPRO Ruby protein stain showing proteins isolated by streptavidin-pulldown. Approximately three hundred embryos transiently expressing each TurboID constructs were subjected to streptavidin-pulldown after biotin incubation and embryo lysis.

Overall, TurboID showed robust biotin labelling with high spatial resolution in zebrafish embryos. These properties rendered it suitable for pursuing *in vivo* proteomic analyses.

### Conditionally stabilised GFP-binding protein (dGBP) is able to target GFP-tagged proteins in zebrafish

Although we were able to achieve proximity-dependent biotin labelling in zebrafish embryos transiently expressing TurboID by mRNA injection, this method requires the direct injection of a large number of newly fertilised embryos in order to obtain sufficient protein for subsequent mass spectrometry sequencing. It is a labour-intensive exercise when potentially analysing multiple POIs, and new genetic constructs must be generated for each POI. In addition, the protein expressed from mRNA injected at the one-cell stage becomes progressively depleted and is present only in trace amounts beyond three days post fertilisation. As such, this methodology is limited to early stage embryos. To circumvent these issues, we envisaged a modular system that would utilise the many existing stable zebrafish lines which express GFP-tagged proteins. Previously, we demonstrated that a GFP-binding peptide (GBP; a 14-kDa nanobody) is able to target a peroxidase (APEX2) to GFP-tagged POIs in both cell culture and zebrafish systems (Ariotti et al., 2015), and can be used for ultrastructural localisation. Based on these findings, we reasoned that genetically fusing TurboID with GBP would target the TurboID-GBP fusion protein to GFP-labelled POIs and/or subcellular compartments in zebrafish, enabling GFP-directed proximity biotinylation *in vivo*. Furthermore, generation of a stable zebrafish line expressing TurboID-GBP would allow delivery of the transgene by a simple genetic cross, circumventing the need for microinjection and enabling continued expression beyond the embryonic stages.

As proof-of-principle, we fused a red fluorescent protein (mRuby2) with the GBP nanobody and transiently expressed it in a transgenic fish line already expressing Cavin1a-Clover. Clover is a GFP derivative recognised by GBP (Shaner et al., 2013), and Cavin1a is an ortholog of caveolae-associated protein 1 in zebrafish (Lo et al., 2015). When expressed at low levels, mRuby2-GBP showed clear colocalisation with Cavin1a-Clover at the plasma membrane in the mRuby2-positive muscle cells (Figure 3A). However, when mRuby-GBP was expressed at higher levels, red fluorescence was observed in the cytoplasm in addition to the plasma membrane, likely due to the saturation of binding between GBP and GFP. This observation raised concerns about the potential of non-specific labelling from unbound TurboID-GBP under these conditions.

**Figure 3.**
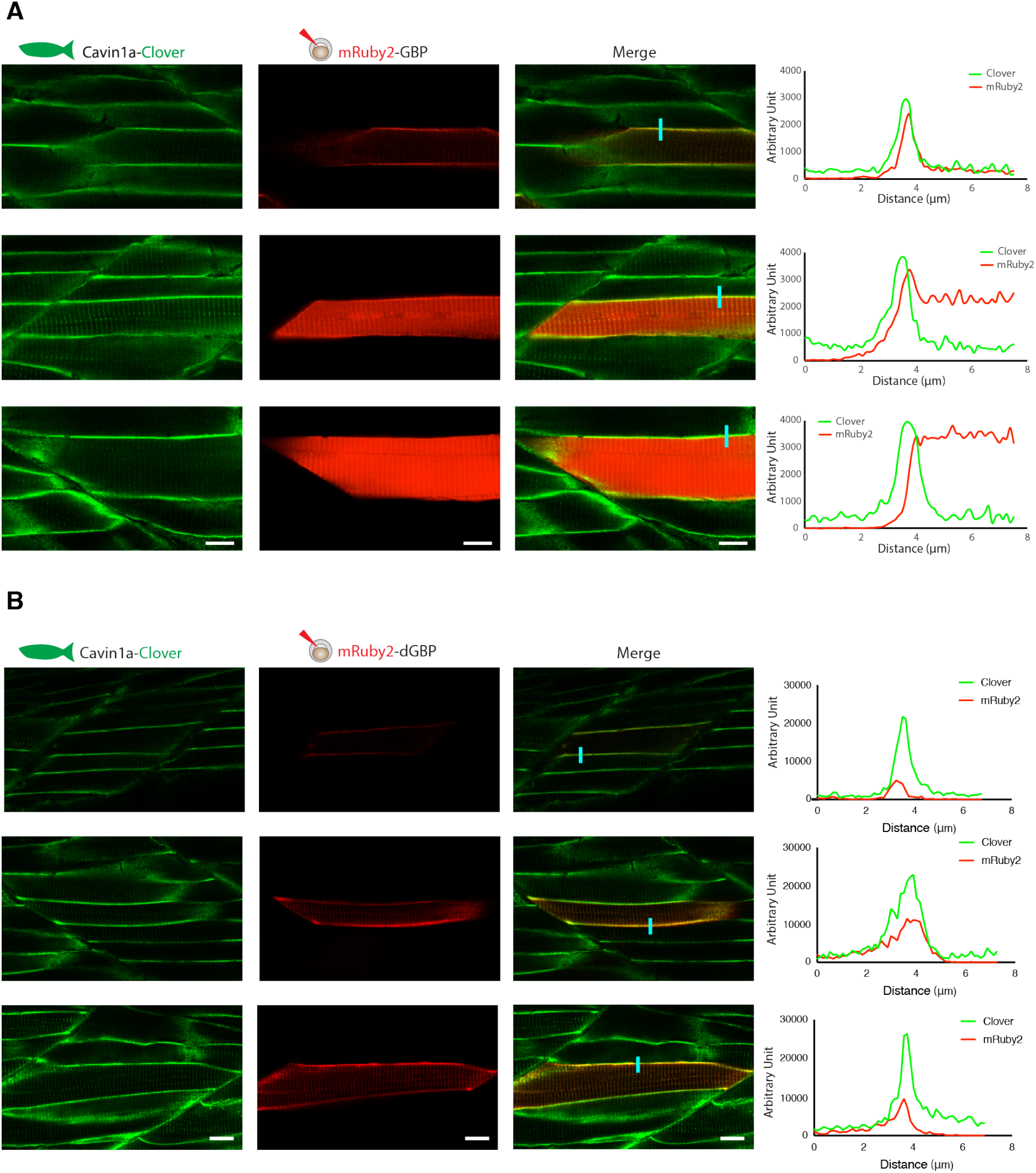
In vivo binding of GFP-nanobody, GBP and dGBP, in stable transgenic zebrafish embryos. (**A** and **B**) Representative images showing the colocalisation between Cavinla-Clover and mRuby2-GBP/dGBP in live zebrafish embryos. Cavinla-Clover zebrafish embryos transiently expressing mRuby2-tagged GBP (**A**) or dGBP (**B**). Injected embryos were imaged at 3 dpf. Line scan (indicated by the blue line) shows the fluorescent intensity of Clover and mRuby2 across the sarcolemma of mRuby2-positive muscle cells. Scale bar denotes 10 μm in both (**A**) and (**B**).

As a solution, we substituted the GBP with a conditionally stabilised GFP-nanobody (destabilised GBP or “dGBP”) that is rapidly degraded unless the GFP-binding site is occupied (Tang et al., 2016; Ariotti et al., 2018). Using this approach, we observed tight association of mRuby2-dGBP and Cavin1a-Clover in all muscle cells regardless of expression level (Figure 3B). We reasoned that a system utilising the conditionally stabilised nanobody would be less likely to result in non-specific biotin labelling within target cells *in vivo*. Furthermore, use of the conditionally stabilised GBP gives potential for modularity, since tissue or cell type specific biotinylation will only occur in cells expressing both GFP-POI and TurboID-dGBP fusion proteins.

### Development of BLITZ; Biotin Labelling in Tagged Zebrafish

We next generated a number of fish lines expressing TurboID-dGBP under the ubiquitous beta actin 2 (*actb2*) promoter (Casadei et al., 2011). To facilitate selection of appropriate transgenic integrations, we added a cytoplasmic red fluorescent protein, mKate2, as a visible reporter upstream of TurboID-dGBP linked by a P2A sequence (Donnelly et al., 2001; Kim et al., 2011). The P2A sequence is a short ribosome-skipping sequence which separates the upstream mKate2 from downstream TurboID-dGBP, reducing the potential interference from the fluorescent protein.

We first tested whether these zebrafish lines were able to catalyse specific biotinylation in tissues expressing GFP. The TurboID-dGBP fish were outcrossed with transgenic lines expressing cytoplasmic GFP in the vasculature (kdrl:EGFP) (Beis et al., 2005) and the motor neurons (MotoN:EGFP) (Punnamoottil et al., 2015) (Figure 4A). Biotinylated proteins were examined in 3 dpf embryos after overnight biotin incubation. In embryos co-expressing ubiquitous TurboID-dGBP and tissuespecific GFP, TurboID-catalysed biotinylation was detected in the intersegmental vessels and the spinal cord motor neurons in the kdrl:EGFP and MotoN:EGFP lines, respectively (Figure 4A). These results demonstrate that our TurboID-dGBP system can produce biotinylation with tissue specificity.

**Figure 4.**
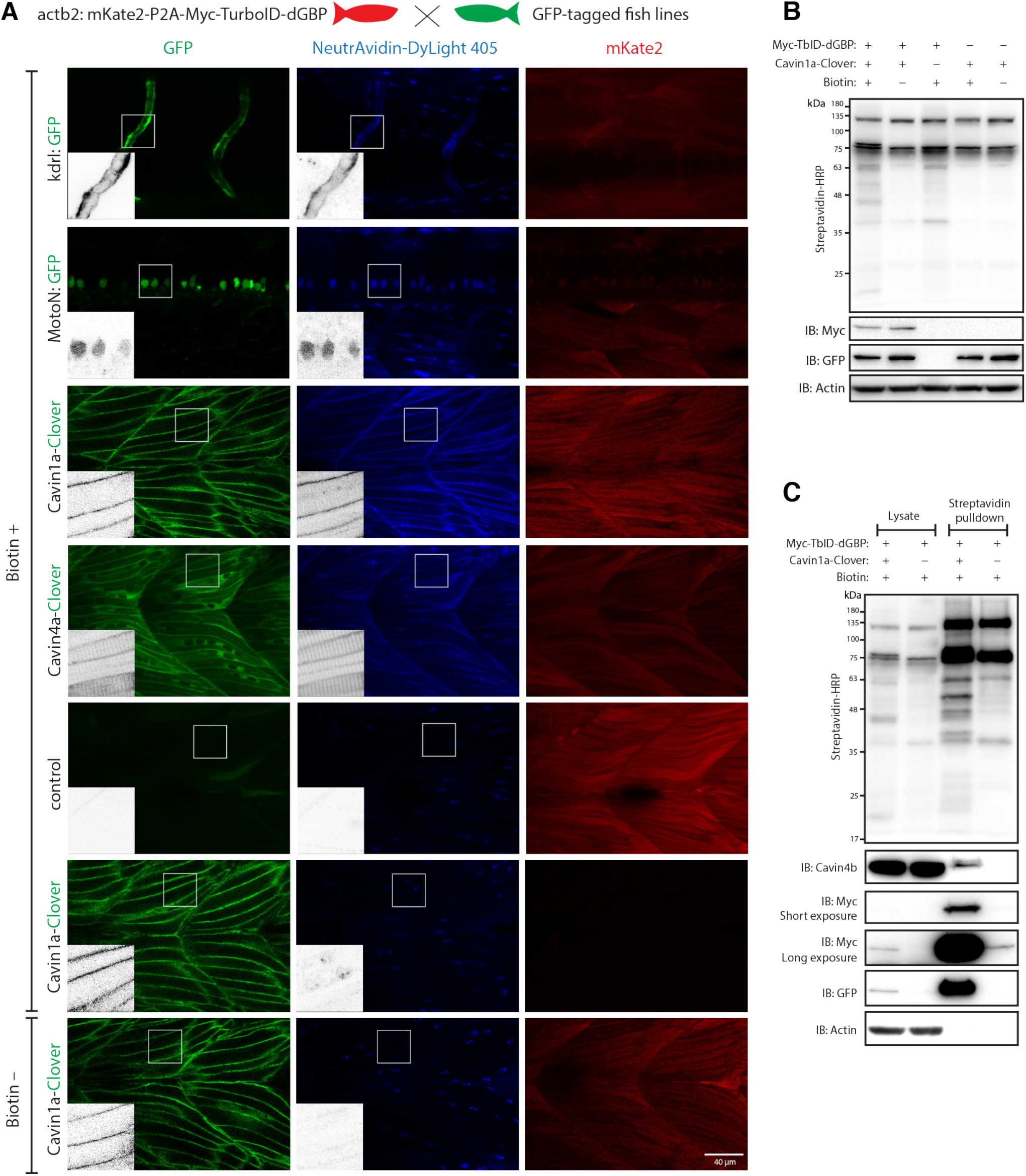
GFP-directed in vivo biotin labelling. (**A**) Representative images of TurboID-dGBP catalysing GFP-dependent biotinylation in transgenic zebrafish embryos at 3 dpf. The TurboID-dGBP line was crossed with different GFP-tagged zebrafish lines: Cavin1a-Clover (plasma membrane), Cavin4a-Clover (sarcolemma and T-tubules), kdrl:eGFP (vasculature), and MotoN:eGFP (motor neurons). After biotin incubation, embryos were fixed, permeabilised and stained with NeutrAvidin to visualise the biotinylated protein. mKate2 is a fluorescent indicator for expression of TurboID-dGBP. Controls were carried out by using siblings from the same clutch without GFP expression and siblings without TurboID expression, as well as omitting biotin incubation. The scale bar denotes 40 μm; n=3. (**B**) Western blot analysis showing the biotinylated proteins in 3 dpf zebrafish embryos from TurboID-dGBP outcrossing with Cavin1a-Clover line. Each sample is a pool of 30 embryos. (**C**) Western blot analysis of fish lysates and streptavidin pulldown with embryos from TurboID-dGBP line outcrossing with Cavin1a-Clover line. Each pulldown sample is a pool of 200 embryos.

To test the biotinylation on a subcellular level, the TurboID-dGBP fish were outcrossed with Cavin1a-Clover and Cavin4a-Clover transgenic fish lines expressing Cavin1a-Clover and Cavin4a-Clover under the control of the muscle specific alpha actin promoter, *acta1a*. Cavin1a and Cavin4a are orthologues of human CAVIN1 and CAVIN4, which are caveola-associated proteins involved in caveolar formation. With the same procedures, we observed clear colocalisation between biotinylated proteins and Clover-tagged cavins in muscle fibres, at the sarcolemma and T-tubules, suggesting our TurboID-dGBP system can produce proximity-dependent biotinylation with subcellular resolution. Without biotin treatment or without the expression of GFP, there was no detectable biotinylation effected by TurboID. Notably, the specificity of GFP-directed biotinylation was not compromised in fish lines expressing higher levels of TurboID-dGBP (Supplementary Figure 3A).

We next visualised the proteins biotinylated by TurboID-dGBP on streptavidin blots (Figure 4B). The two prominent bands representing endogenously biotinylated proteins were again observed in embryos carrying both TurboID-dGBP and Cavin1a-clover; omitting the biotin supplement resulted in no exogenous biotinylation. Intriguingly, in the absence of Cavin1a-Clover, a weak biotinylation was still observed in the embryos carrying only the TurboID-dGBP transgene, despite the level of TurboID-dGBP being undetectable on anti-Myc immunoblot. This background labelling is likely caused by TurboID-dGBP en route to proteasomal degradation. Using streptavidin affinity pulldown, biotinylated proteins were isolated from total fish lysates and endogenous Cavin4b, a known Cavin1 interactor (Bastiani et al., 2009), was detected in the streptavidin pulldown in addition to Cavin1a-GFP and TurboID-dGBP (Figure 4C). Note that a trace of TurboID-dGBP was detected in the streptavidin pulldown in the absence of GFP target with long exposure, which accounts for the weak background biotinylation in embryos expressing only TurboID-dGBP.

### A comprehensive cavin proteome in skeletal muscle generated by TurboID-dGBP

Finally, we employed our TurboID-dGBP system to map the proteomes associated with Cavin1 and 4 in zebrafish skeletal muscle. We crossed the TurboID-dGBP fish with fish lines stably expressing Cavin1a-Clover, Cavin4a-Clover and Cavin4b-Clover in muscle. TurboID-dGBP and cavin-Clover co-expressing embryos were selected at 2 dpf for subsequent biotin labelling and the biotinylated proteins were analysed by liquid chromatography coupled to tandem mass spectrometry (nanoHPLC/MS MS/MS).

In total, we identified 277, 178 and 137 proteins in the Cavin1a, Cavin4a and Cavin4b samples, respectively, with high confidence (at least one high confidence peptide detected in the sample) (Supplementary file 1). An enrichment score (ES) defined as 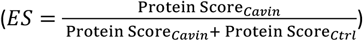, was calculated for each identified protein. Proteins only identified in the Cavin group will have an enrichment score of 1, while endogenous biotinylated proteins and non-specific binding proteins will tend to have an enrichment score around 0.5 (Figure 5A-C). By using ES > 0.9 as a cut-off, 83, 82, and 39 proteins were found enriched in Cavin1a, Cavin4a and Cavin4b samples, respectively (Figure 5D; Table 1; Supplementary file 2). Among the proteins detected in the Cavin1a sample, known Cavin1 interactors, such as endogenous caveolins and cavins, were identified with an ES of 1 (Figure 5A, highlighted in green). Dystrophin (Dmd), a protein associated with caveolae (Song et al., 1996; Doyle et al., 2000), was also highly enriched in the Cavin1a group (ES of 0.99). An ortholog of Pacsin3, a caveola-associated protein required for muscle caveolar formation, (Seemann et al., 2017) was also enriched in the Cavin4b group (ES of 1.0). These interactors were undetectable in all control samples, demonstrating the high accuracy of the BLITZ system. Individual hits and general properties of putative interacting proteins are further discussed below.

**Figure 5.**
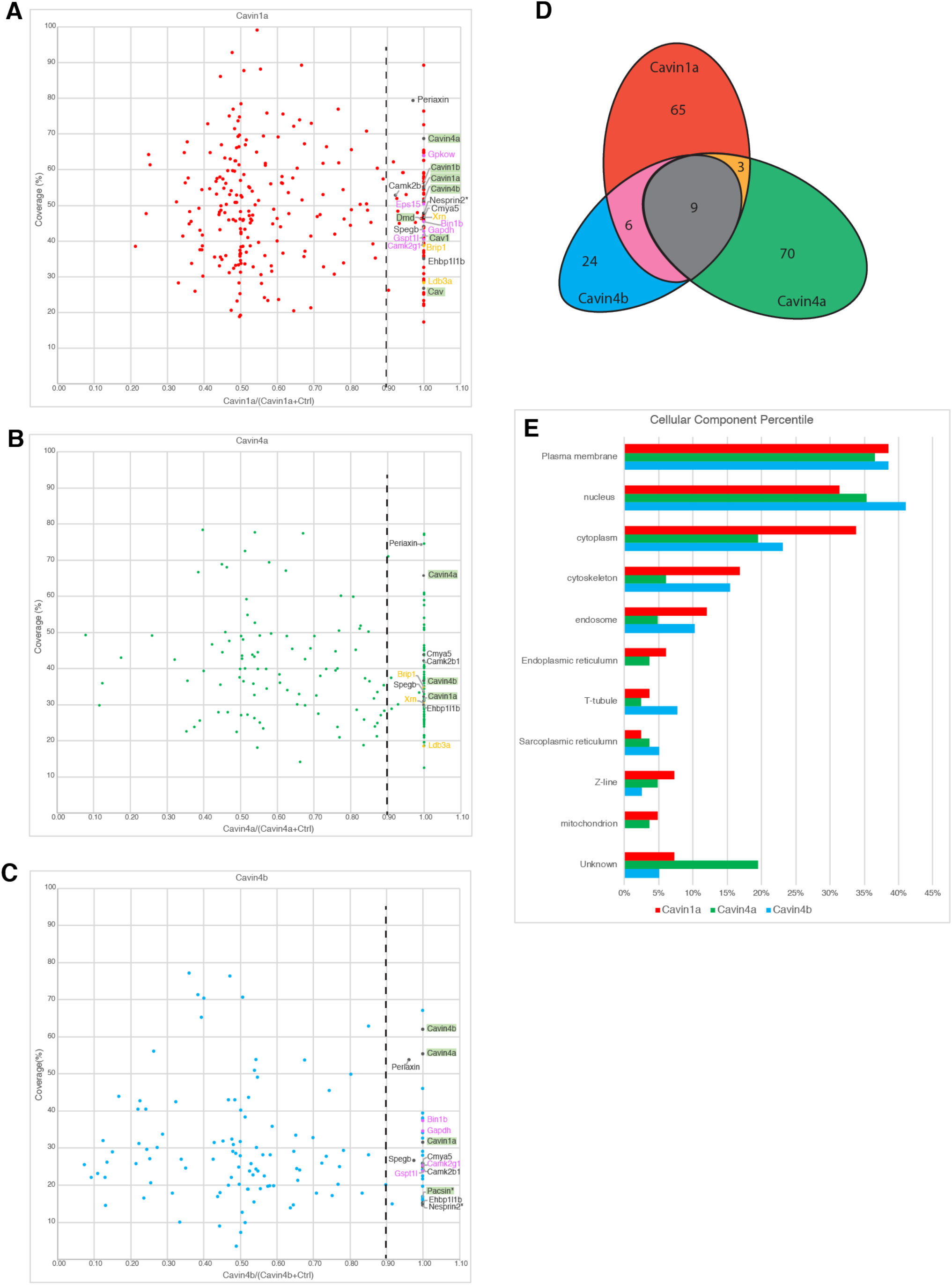
Proteomes identified by BLITZ system in Clover-tagged cavin zebrafish. (**A**, **B** and **C**) Scatter plot showing proteins identified by BLITZ with Cavin1a-Clover (**A**), Cavin4a-Clover (**B**) and Cavin4b-Clover (**C**) lines. The Y axis represents protein coverage and the X axis represents the enrichment score. Known interactors or proteins in close proximity were highlighted in green. Protein name in black indicates the proteins were identified in all cavin samples; orange indicates the proteins were identified in both Cavin1a and Cavin4a samples; pink indicates proteins identified in both Cavin1a and Cavin4b samples. (**D**) Venn diagram showing the overlap of identified proteins in Cavin1a, Cavin4a and Cavin4b samples. (**E**) Bar graph showing the distribution of proteins at subcellular level. The cellular component information was curated from Uniport database and literature.

**Table 1.**
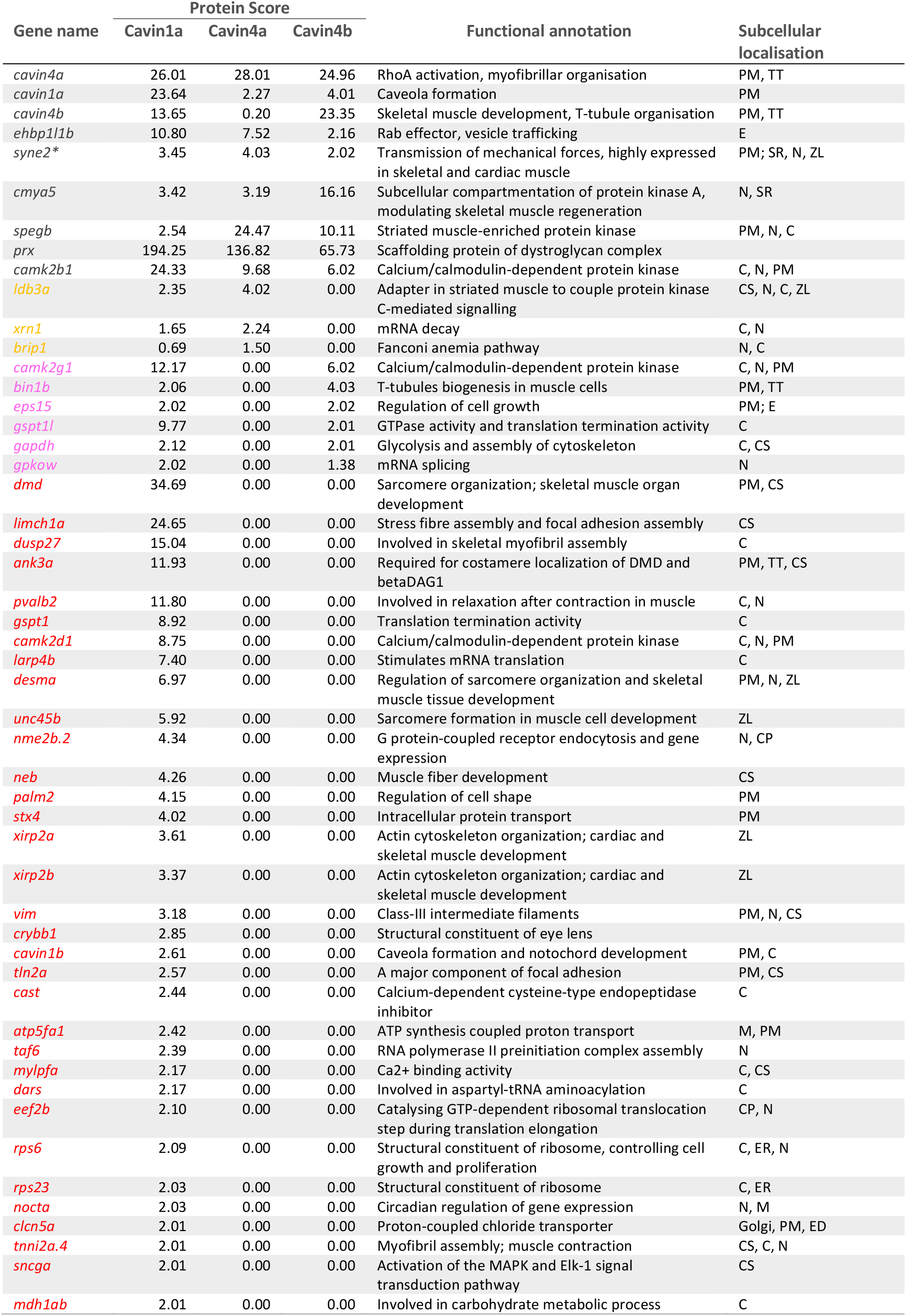
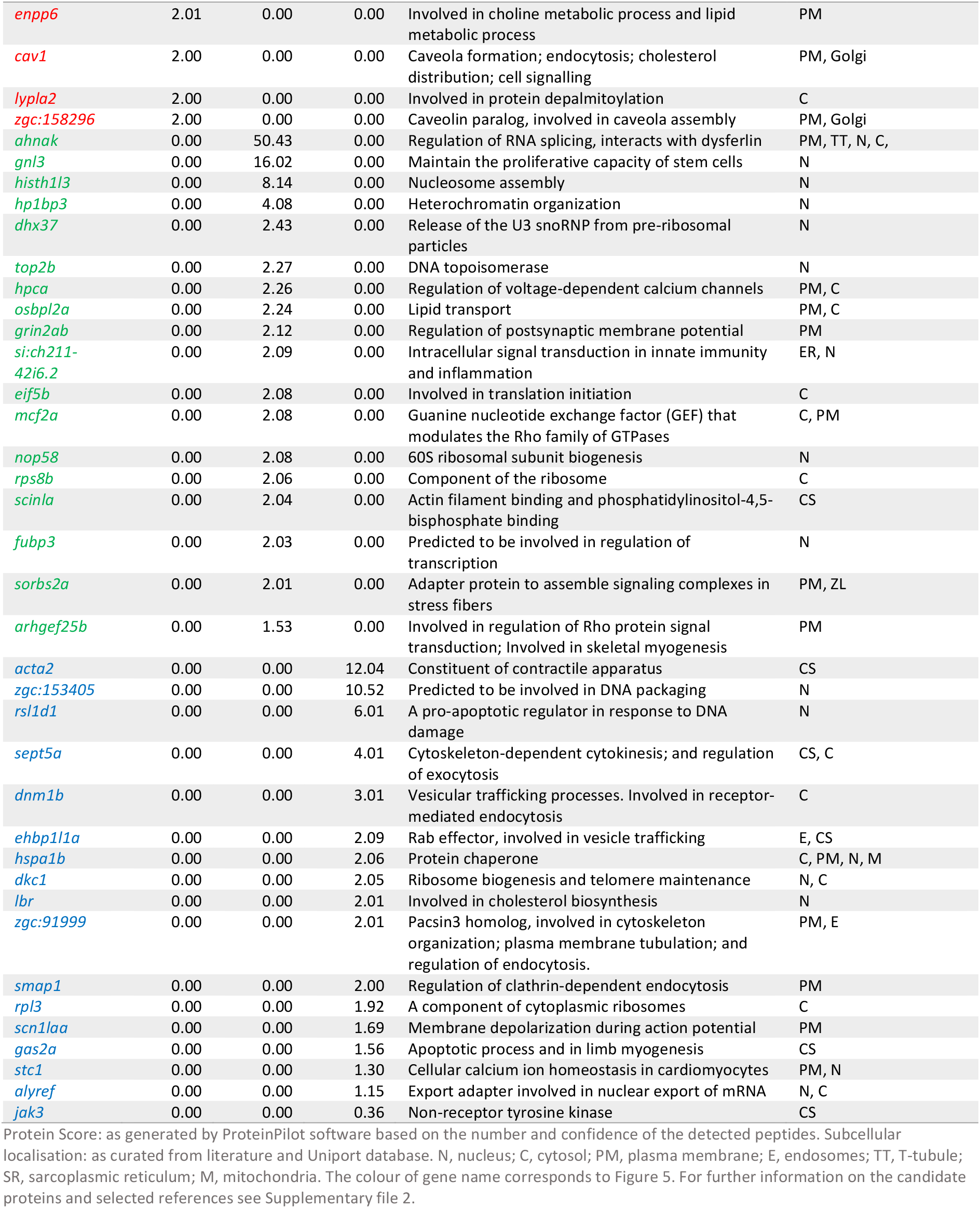
Top cavin interactor candidates from Figure 5

## Discussion

### Advantages of BLITZ in proteomic mapping

In this study, we have developed BLITZ (Biotin Labelling In Tagged Zebrafish): a modular system for *in vivo* proteomic mapping (Figure 6). This system utilises the advantages of BioID at capturing weak or transient interactions in living cells, but extends its application to an *in vivo* setting, enabling interactome investigation at specific developmental stages under physiological conditions, and potentially in disease models. The system also has several advantages over conventional BioID methods for studying the proteome and interactome.

**Figure 6.**
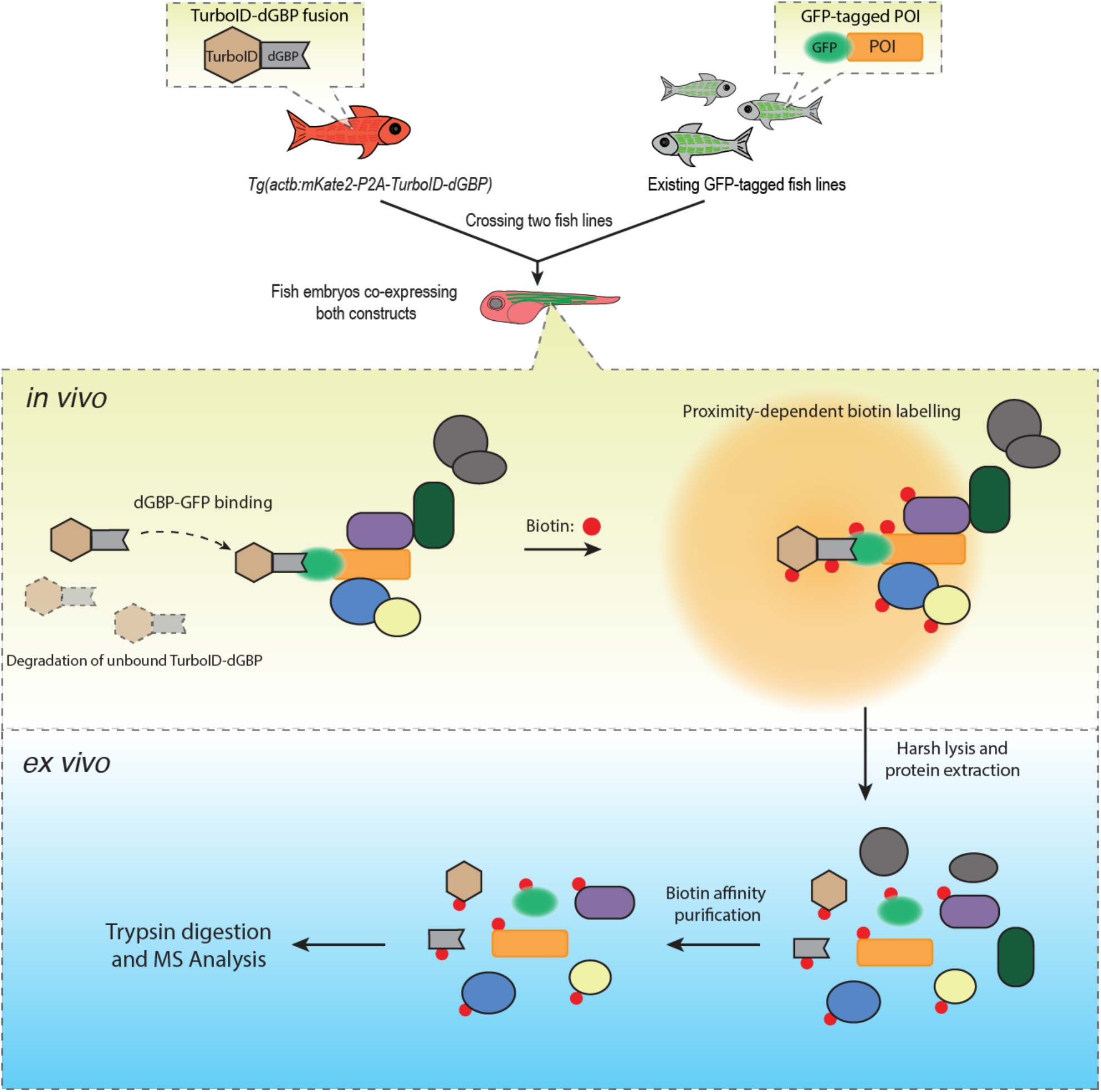
A schematic overview of the BLITZ system. The TurboID-dGBP lines can be crossed with existing GFP-tagged lines. In the embryos carrying both transgenes, the binding between dGBP and GFP stabilise TurboID-dGBP, which leads to proximity biotinylation around the GFP-tagged POIs. The unbound TurboID-dGBP will be rapidly degraded by the ubiquitin proteasome system, which minimises non-specific labelling when dGBP-GFP binding saturates, as well as achieving tissue specificity by averting labelling in cells/tissues that do not express GFP. The biotin labelled proteins can be isolated by biotin affinity purification and identified by MS analysis.

Firstly, BLITZ does not require extensive molecular biology steps to produce numerous expression constructs, or laborious embryonic manipulation. It instead relies on simple crossing of a TurboID-dGBP line with an existing GFP-tagged fish line of choice; a plethora of such lines currently exist in stock centres globally and, with the advent of nuclease directed genome editing, this number is rapidly increasing. Secondly, BLITZ enables cell- and tissue-specific proteomic studies, since the stability of TurboID-dGBP is dependent on its binding to GFP targets. Non-specific biotin labelling in tissues that do not express GFP-tagged constructs is avoided. Thirdly, our TurboID-dGBP system has the potential to be used with knock-in fish lines carrying a GFP fusion protein at the endogenous locus of a POI. This will enable the application of BioID to study proteomic associations with endogenous proteins, which, to our knowledge, has not been achieved by conventional BioID methods. Finally, the modularity of the BLITZ system could be advantageous for use in established tissue culture systems using existing GFP expression vectors or/and knock-in cell lines, as well as extended to other organisms.

The use of the BLITZ system also comes with some caveats. Unlike the traditional BioID approach using a direct fusion of the biotin ligase with the bait protein, our system targets TurboID to the POI through the binding of dGBP nanobody to GFP. In this case, the indirect binding increases the physical distance between biotin ligase and the POI, which could potentially enlarge the effective labelling radius and include more non-interacting neighbouring proteins. However, we have previously shown that the use of a GFP-directed nanobody to target a genetically encoded peroxidase (APEX2) for protein localisation does not appear to compromise the fidelity of labelling: APEX2 staining was rarely observed beyond 25 nm from the site of POI (Ariotti et al., 2015). It is also possible that the binding of the biotin ligase-nanobody with the GFP-tagged POI could perturb the localisation of the POI, either by masking interacting surfaces or simply due to the larger size of a complex. For this reason, we routinely examine the distribution of the GFP-tagged POI both with and without biotin ligase-dGBP expression as well as the distribution of biotinylated proteins using fluorescent neutravidin staining. Since our *in vivo* system is based on the simple crossing of heterozygous transgenic lines, every new clutch contains offspring with every possible combination of alleles, and the appropriate internal controls can be sorted by fluorescence.

### Application of BLITZ to the identification of cavin-association networks in muscle

Cavin family proteins are key components of the caveolar coat complex associated with the inner leaflet of the plasma membrane. Cavin1 is present in all tissues and is essential for caveolar formation and function. Cavin2, 3 and 4 show more restricted tissue distributions with Cavin4 being specific to skeletal and cardiac muscle (reviewed in Parton et al., 2018). In the zebrafish, Cavin1 and 4 are each duplicated such that four loci exist; Cavin1a/b and Cavin4a/b. Cavin1a and b show spatially distinct expression patterns with Cavin1b being largely restricted to the developing notochord whereas Cavin1a, 4a and 4b are all highly expressed in skeletal muscle (Hill et al., 2008; Lo et al., 2015; Housley et al., 2016). In this study we used the BLITZ system to identify putative interactors for all three skeletal muscle cavins, and identified sets of putative interactors both unique and common to all three proteins.

The majority of proteins identified for all cavin proteomes were muscle-enriched factors and plasma membrane proteins. We also saw a specific enrichment of known caveola-associated proteins consistent with initial expectations (Table 1; Supplementary File 2). However, the cavin proteomes also contained a disproportionate number of nuclear proteins, such as Brip1, Xrn1, Taf6. In cultured cells, cavins have been shown to be released from the plasma membrane in response to external stimuli (such as mechanical stress) and are able to bind intracellular targets in variety of subcellular locations to regulate processes such as ribosomal RNA transcription and apoptosis (Liu and Pilch, 2016; McMahon et al., 2019). In addition, in the absence of Cavin1, in knockout mouse muscle, Cavin4 has been shown to localise predominantly to the nucleus rather than the sarcolemma (Lo et al., 2015).

What processes might Cavin1 and Cavin4 be regulating? We know that loss of Cavin1 causes lipodystrophy and muscular dystrophy in humans. Patient and animal muscle shows hypertrophied muscle fibres (Hayashi et al., 2009; Rajab et al., 2010; Ding et al., 2017). Cavin4 mutations have been described in dilated cardiomyopathy patients and there is evidence that Cavin4 recruits ERK in cardiomyocytes (Rodriguez et al., 2011; Ogata et al., 2014). Thus, there is supporting data for the positive regulation of hypertrophy in skeletal muscle fibres by Cavin1, and in cardiomyocytes by Cavin4. The cavin proteome showed an enrichment of protein kinases, such as calcium/calmodulin-dependent protein kinase II (CaMKII). CaMKII regulates Ca^2+^ signalling and plays an important role in the development of cardiac hypertrophy through the ERK signalling pathway (Illario et al., 2003; Cipolletta et al., 2010; Cipolletta et al., 2015). The activation of CaMKII can be induced by exercise in skeletal muscle, with the activation level proportional to the intensity of exercise (Rose et al., 2006).

In this study, BLITZ revealed several putative cavin interactors that have also been shown to be involved in cardiomyopathies and/or skeletal myopathies, including the membrane protein Dystrophin (Deconinck and Dan, 2007), the intermediate filament protein Desmin (Hnia et al., 2015), and the triad associated proteins Bin1 (Nicot et al., 2007), Cypher/ZASP (Selcen and Engel, 2005) and SPEG (Agrawal et al., 2014). Genetic ablation of zebrafish Cavin4b causes aberrant T-tubules in skeletal muscle (Housley et al., 2016). It is possible that Cavin4 may be involved in T-tubule formation through interaction with triad associated proteins, such as Bin1.

Overall, our BLITZ system enables the *in vivo* identification of protein interactors in a cell- and tissue-specific manner, with high precision. We demonstrated the applicability of this approach in diverse cell types including neurons and vascular endothelial cells and applied the BLITZ system to identify factors associated with cavin family proteins in differentiated skeletal muscle. BLITZ provides a versatile and valuable tool for proteomic discovery using the zebrafish model, but also has the potential for application in other *in vivo* contexts that to date have been challenging or intractable.

## Materials and methods

**Table 2.**
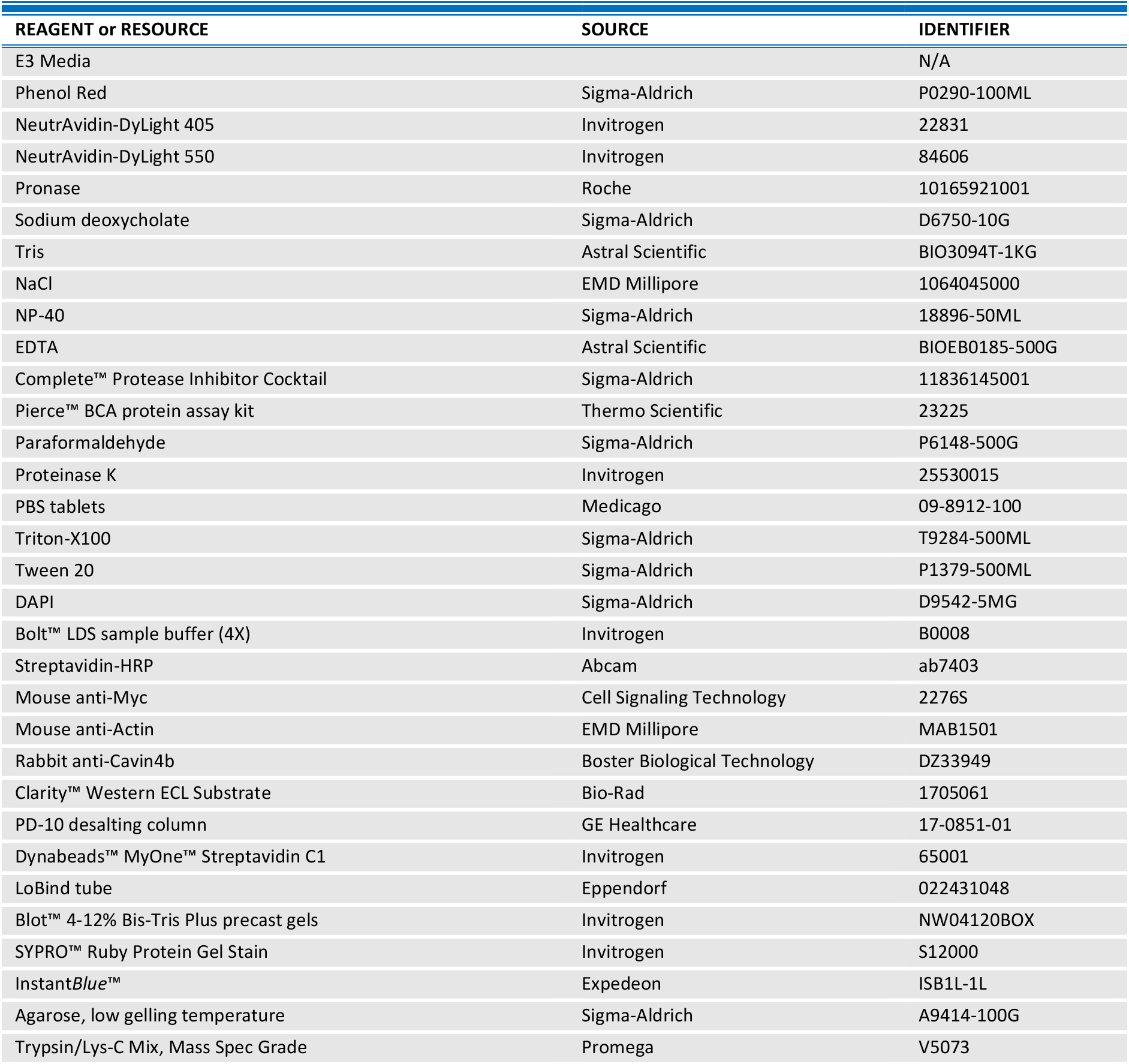

### Zebrafish husbandry

Zebrafish were raised and maintained according to institutional guidelines (Techniplast recirculating system, 14-h light/10-h dark cycle, University of Queensland, UQ). Adults (90 dpf above) were housed in 3 or 8 L tanks with flow at 28.5 °C and embryos up to 5 dpf were housed in 8 cm Petri dishes in standard E3 media (5 mM NaCl, 0.17 mM KCl, 0.33 mM CaCl2 and MgSO4) at 28°C (incubated in the dark) (Westerfield, 2007). All experiments were approved by the University of Queensland Animal Ethics Committee. The following zebrafish strains were used in this study: wild-type (TAB), an AB/TU line generated in UQBR Aquatics (UQ Biological Resources), *Tg(acta1:cavin1a-Clover), Tg(acta1:cavin4a-Clover), Tg(acta1:Cavin4b-Clover), Tg(actb2:mKate2-P2A-TurboID-dGBP), Tg(kdrl:eGFP) and Tg(MotoN:GFP)*. The developmental stages of zebrafish used in experiments are prior to specific sex determination. All zebrafish used in experiment were healthy, not involved in previous procedures and drug or test naïve.

### DNA constructs and transgenic fish lines

The protein sequence of TurboID and MiniTurbo was constructed according to Branon et al., 2018 while the protein sequence of BASU was designed according to Ramanathan et al., 2018. The coding sequences of TurboID, MiniTurbo and BASU were ordered from IDT as gene fragment with codon optimised for zebrafish expression (https://sg.idtdna.com). The expression of biotin ligases was driven by a ubiquitous promoter of *actb2* (Higashijima et al., 1997; Casadei et al., 2011). A red fluorescent reporter, mKate2, was indirectly linked into the N-terminus of biotin ligase through a self-cleaving P2A sequence (Shcherbo et al., 2009; Kim et al., 2011). Promoter, fluorescent report and biotin ligase were cloned into destination vector using Gateway cloning system. All fish lines were generated by using Tol2kit system according to established methods (Kawakami, 2004; Kwan et al., 2007). In brief, plasmid constructs for generating transgenic lines were coinjected with tol2 mRNA into one-cell-stage WT zebrafish embryos (Nusslein-Volhard and Dahm, 2002). Injected F_0_s were raised and screened for founders producing positive F_1_s with Mendelian frequencies, indicative of single genomic integration. Positive F_1_s grown to reproductive age were used for our experiments. Stable lines were maintained as heterozygotes. All stable lines used are given in table below.

**Table 3.**

| Transgenic Fish lines | Details | reference |
| --- | --- | --- |
| TurboID-dGBP | *Tg(actb2:mKate2-P2A-TurboID-dGBP)* | generated in this paper |
| Cavin1a-Clover | *Tg(acta1:Cavin1a-Clover)* | generated in this paper |
| Cavin4a-Clover | *Tg(acta1:Cavin4a-Clover)* | generated in this paper |
| Cavin4b-Clover | *Tg(acta1:Cavin4b-Clover)* | generated in this paper |
| Kdrl:GFP | *Tg(kdrl:EGFP)* | (Beis et al., 2005) |
| MotoN:GFP | *Tg(miR218:EGFP)* | (Punnamoottil et al., 2015) |
| Constructs | Addgene identifier |  |
| pT3TS-BASU-EGFP (constructs for RNA) | Pending |  |
| pT3TS-TurboID-EGFP (constructs for RNA) | Pending |  |
| pT3TS-miniTurbo-EGFP (constructs for RNA) | Pending |  |
| pT3TS-TurboID-CaaX (constructs for RNA) | Pending |  |
| pT3TS-NLS-TurboID (constructs for RNA) | Pending |  |
| pT3TS-CD44b-TurboID (constructs for RNA) | Pending |  |
| pT3TS-Cavin4b-TurboID (constructs for RNA) | Pending |  |
| Cavin1a-Clover (construct for generating stable fish line) | Pending |  |
| Cavin4a-Clover (construct for generating stable fish line) | Pending |  |
| Cavin4b-Clover (construct for generating stable fish line) | Pending |  |
| mRuby2-GBP(DNA construct) | Pending |  |
| mRuby2-dGBP(DNA construct) | Pending |  |
| mKate2-P2A-TurboID-dGBP (constructs for generating stable fish line) | Pending |  |

### Transient expression by DNA/RNA microinjection

DNA plasmid and RNA transcript for injection were diluted to final concentration of 30ng/μl and 200ng/μl, respectively, with addition of Phenol Red (Sigma-Aldrich) as injection tracer. A bolus of approximately 1/5 of the total cell diameter was injected into each embryo. For DNA injection, the bolus was injected into the cell of embryos at single cell stage (5 to 25 min-post-fertilisation). For RNA injection, the bolus was injected into the yolk of the embryos up until 2-ell stage. The RNA transcript was synthesised by mMESSAGE mMACHINE™ T3 (Invitrogen) according to manufacturer’s instruction. The RNA transcripts were tagged with poly(A) tail using Poly(A) Tailing Kit (Invitrogen) to extend the stability of mRNA in zebrafish embryos.

### *In vivo* biotin labelling

Embryos at indicated developmental stage were incubated in the E3 media supplemented with 500 μM biotin for 18 h to initiate biotinylation *in vivo*. For embryos before hatching, a dechorionation step was carried out by using Pronase (Roche, 100 μg/ml in E3 media for 40 min at 28°C) prior to the biotin incubation. After biotin incubation, embryos were washed for 40 min with two changes of standard E3 media to remove unincorporated biotin before subsequent immunostaining or protein extraction.

### Zebrafish embryos protein extraction

Fish embryos after *in vivo* biotin labelling were deyolked by mechanical disruption through a 200 μl pipette tips in calcium-free Ringer’s solution followed by two changes of solution at 4 °C. The deyolked embryos was lysed by brief sonication in RIPA buffer (50 mM Tris-HCl, pH 7.5; 150 mM NaCl; 1% NP-40; 0.1% SDS; 5 mM EDTA; 0.5% Na-deoxycholate,) with freshly added cOmplele™ Protease Inhibitor. Lysates were further solubilised at 4 °C with rotation for 30 min. Insoluble material was removed by centrifugation at 14,000 ×g for 10 min at 4 °C, and supernatant were collected for BCA protein assay determining protein concentration. For Western blot analysis, 25 fish embryos per group were used for protein extraction, whilst, for streptavidin affinity purification, approximately 350 embryos were used for each group.

### Western blotting

Western blot analysis was performed largely as described previously Lo *et al*., 2015. Briefly, zebrafish samples from protein extraction were mixed with NuPAGE™ LDS sample buffer (4X) and 10 mM DTT. Protein samples were analysed by Western blotting with following antibodies: mouse anti-Myc (dilution 1:2000), mouse anti-Actin (dilution 1:5000), rabbit anti-Cavin4b (dilution 1:2000), anti-mouse and anti-rabbit HRP-conjugated antibodies (dilution 1:5000), streptavidin-HRP (dilution 1:5000). ECL blotting reagent was used to visualise HRP and chemiluminescent signal was detected using the ChemiDoc MP system (BioRad) as per the manufacture’s instruction.

### Streptavidin beads pulldowns

Fresh embryo protein extracts (4 mg in 2.5 ml RIPA buffer) was passed through PD-10 desalting column (GE Healthcare) to remove excess free biotin using the gravity protocol according to manufacturer’s instruction. Protein extracts were then mixed with Dynabeads MyOne Streptavidin C1 (Invitrogen) from 200 μl bead slurry that were pre-washed with RIPA buffer, and incubated on a rotor wheel at 4 °C overnight (16 h). The next day, the beads were separated from the protein extracts on a magnetic rack and transferred to a new 2 ml LoBind tube (Eppendorf). The beads were washed with 1 ml of each following solution: twice with RIPA buffer, once with 2% SDS in 50 mM Tris-HCl pH7.5, once with 2 M urea in 10 mM Tris-HCl pH8.0 and twice again in RIPA buffer without cOmplete Protease Inhibitor. Washed beads were boiled in 60 ul of 2X Blot LDS sample buffer (4X diluted to 2X with RIPA buffer) containing 2 mM biotin and 20 mM DTT at 95 °C for 10 min with 10 s vortex after first 5 min boiling. 5 ul of the pulldown samples was used for immunoblots, while the 50 ul of the samples were used for SDS-PAGE with SYPRO Ruby (Invitrogen) or InstantBlue protein gel stain ().

### Immunostaining and confocal microscopy

Fish embryos after *in vivo* biotin labelling were fixed in 4% paraformaldehyde (PFA) overnight at 4 °C. After fixation, embryos were permeabilised by proteinase K (10 μg/ml, 10 min for embryos at 2 dpf, or 15 min for embryos at 3 dpf), and fixed again with 4% PFA for 15 min. Embryos were washes with PBS-Tween 20 (0.1%) and blocked in PBS with 0.3% Triton X-100 and 4% BSA for 3 h at room temperature. Staining was performed in blocking buffer with NeutrAvidin-DyLight (1:500 dilution) overnight at 4 °C followed by 4 washes with PBS 0.3% Trition X-100. For nuclear staining, embryos were stained with DAPI for 10 min followed by 3 washes with PBS 0.3% Trition X-100.

Confocal imaging was performed on Zeiss 710 meta upright confocal microscopes. Zebrafish embryos were mounted in 1% low melting point agarose in embryos media (Westerfield, 2007) on a standard 8 cm petri dish. Objectives used were Zeiss water immersion x40 N/A 1.0 (catalogue number 420762). For live embryo imaging, embryos were anaesthetised in 2.5 mM tricaine prior to imaging.

### Sample preparation for mass spectrometry

For in-gel digestion, the streptavidin pulldown samples were separated by SDS-PAGE on a 4-12% precast gel (Blot™ Bis-Tris Plus, Invitrogen) and then stained with Instant*Blue* (Expedeon). The whole lane was excised from the gel and future cut into approximate 3 x 1 x 2 mm^3^ slices (L x W x H) and each slice were placed into a separate LoBind tubes (Eppendorf) for destaining. The gels were destained by adding 500 μl of 100 mM ammonium bicarbonate/acetonitrile (1:1, vol/vol) and incubated with occasional vortexing for 30 min. The ammonium bicarbonate/acetonitrile buffer was removed, and the gel pieces dried for 15 min by the addition of 200 μl of acetonitrile. The acetonitrile was removed and another 200 μl aliquot was added and left for 15-30 min. The acetonitrile was removed in preparation for trypsin digestion.

After destain, the gel pieces were covered with 200 μl of 20 ng/μl of sequence grade trypsin/Lys-C (Promega) in 50 mM ammonium bicarbonate pH8 buffer. The gel pieces were left for 1 h and if required a further 100 μl of trypsin/Lys-C solution was added to cover the gel pieces. The samples were placed in an incubator at 37 °C overnight. The trypsin solution was transferred from each sample and placed in a clean Eppendorf tube. 200 μl of 5% formic acid/acetonitrile (3:1, vol/vol) was added to each tube and incubated for 15 min at room temperature in a shaker. The supernatant was placed into the pre-cleaned Eppendorf tubes, together with the trypsin solution for each sample and dried down in a vacuum centrifuge.

For HPLC/MS MS/MS analysis, 12 μl of 1.0% (vol/vol) trifluoroacetic acid in water was added to the tube, which was vortexed and incubated for 2 min in the sonication bath and then centrifuged for 1 min at 6,700 xg (10,000 rpm) and finally, transferred to an autosampler vial for analysis.

### Lipid chromatography and mass spectrometry

The tryptic peptide extracts were analysed by nanoHPLC/MS MS/MS on an Eksigent ekspert™ nanoLC 400 system (SCIEX) coupled to a Triple TOF 6600 mass spectrometer (SCIEX) equipped with PicoView nanoflow (New Objective) ion source. 5 μl of each extract was injected onto a 5 mm x 300 μm, C18 3 μm trap column (SGE, Australia) for 5 min at 10 μL/min. The trapped tryptic peptide extracts were then washed onto the analytical 75 μm x 150 mm ChromXP C18 CL 3 μm column (SCIEX) at 400 nl/min and a column temperature of 45 °C. Linear gradients of 2-40% solvent B over 60 min at 400 nl/min flowrate, followed by a steeper gradient from 40% to 90% solvent B in 5 min, then 90% solvent B for 5 min, were used for peptide elution. The gradient was then returned to 2% solvent B for equilibration prior to the next sample injection. Solvent A consisted of 0.1% formic acid in water and solvent B contained 0.1% formic acid in acetonitrile. The ionspray voltage was set to 2600V, declustering potential (DP) 80V, curtain gas flow 30, nebuliser gas 1 (GS1) 30, interface heater at 150°C. The mass spectrometer acquired 50 ms full scan TOF-MS data followed by up to 30 100 ms full scan product ion data, with a rolling collision energy, in an Information Dependant Acquisition, IDA, mode for protein identification and peptide library production. Full scan TOFMS data was acquired over the mass range 350-1800 and for product ion ms/ms 100-1500. Ions observed in the TOF-MS scan exceeding a threshold of 200 counts and a charge state of +2 to +5 were set to trigger the acquisition of product ion, ms/ms spectra of the resultant 30 most intense ions.

### MS Data analysis and GO annotation

The data was acquired and processed using Analyst TF 1.7 software (SCIEX). Protein identification was carried out using ProteinPilot™ software v5.0 (SCIEX) with Paragon™ database search algorithm. MS/MS spectra were searched against all zebrafish proteins annotated at UniProt database. The peptide evidence from the search was assembled by Pro Group™ Algorithm to generate a list of protein identification with Protein Score (ProtScore). The ProtScore was calculated based on the number and the confidence of identified peptides. The peptide identifications could only contribute to the Unused ProtScore of a protein to the extent that their spectra have not already been used to justify an already assigned more confident protein. Proteins were only reported as detected if they had sufficient Unused ProtScore.

The ProScore of each protein were firstly normalised to the ProtScore of an endogenous biotinylated protein, propionyl-CoA carboxiylase subunit alpha, detected within the group. The enrichment score (ES) was calculated by dividing ProtScore of each protein in cavin group by the combined ProtScore of that protein in both cavin and control groups.

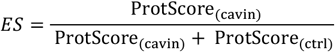

Therefore, a protein that is only identified in cavin group will have an enrichment score of 1, while endogenous biotinylated proteins and non-specific binder would likely have an enrichment score around 0.5.

The GO annotation for our identified proteins was carried out using DAVID 6.8 (Huang da et al., 2009b, a). For zebrafish proteins without annotation information, manually annotation was performed by searching their human or mouse homologs.

## Acknowledgements

We would like to thank Dr. Anne Lagendijk and Dr. Jean Giacomotto for *Tg(kdrl:EGFP)* and *Tg(MotoN:EGFP)* fish lines, respectively. Confocal microscopy was performed at the Australian Cancer Research Foundation (ACRF)/Institute for Molecular Bioscience (IMB) Dynamic Imaging Facility for Cancer Biology with funding from the ACRF. MS spectrometry was performed at the IMB Mass Spectrometry Facility, the University of Queensland. This work was supported by fellowship and grants from the National Health and Medical Research Council of Australia (to R.G. P., grant numbers 569542 and 1045092; to R.G.P., grant number APP1044041; and to T.E. H. and R.G.P., grant number APP1099251), the Australian Research Council (to T.E.H. and P.G.P., grant number DP200102559) as well as by the Australian Research Council Centre of Excellence in Convergent Bio-Nano Science and Technology (to R.G.P., grant number CE140100036).

## Supplementary Figures

**Figure S1.**
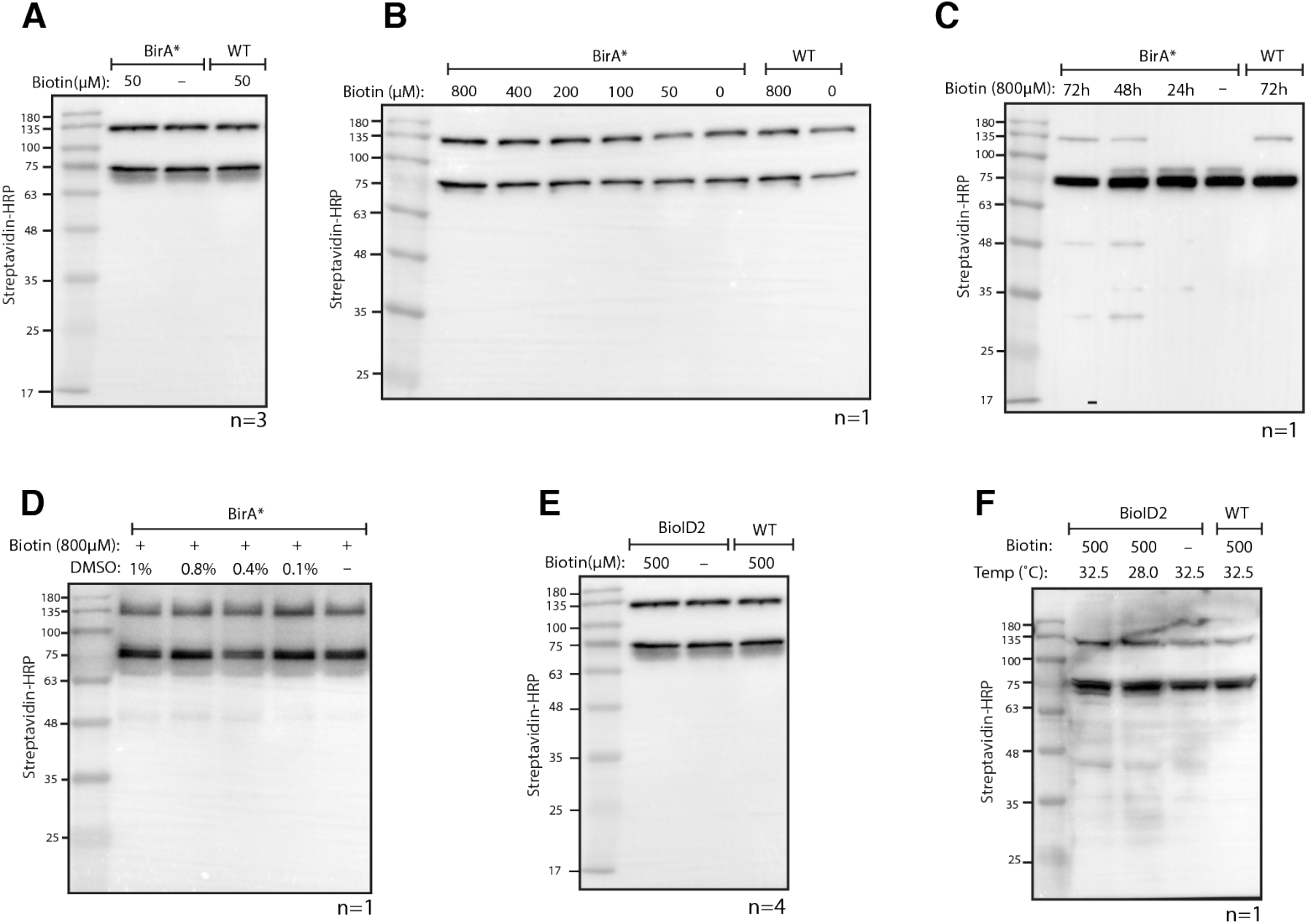
Testing BioID and BioID2 in transgenic zebrafish. (**A**) Streptavidin-HRP blot showing biotinylated proteins in BioID (BirA*) transgenic zebrafish embryos. Zebrafish embryos were incubated with 50 μM biotin for 18 h prior to protein extraction and Western blotting. (**B**, **C** and **D**) Streptavidin-HRP blots showing biotinylated proteins in zebrafish embryos with (**B**) increased biotin concentration, (**C**) increased biotin incubation time and (**D**) addition of DMSO. (**E** and **F**) Streptavidin-HRP blots showing biotinylated proteins in BioID2 transgenic zebrafish embryos under (**E**) standard incubation temperature (28 °C) and (**F**) elevated temperature (32.5 °C) during biotin incubation. Each sample is a pool of 30 embryos at 3 dpf. The two prominent protein bands present in all samples were endogenous biotinylated proteins.

**Figure S2.**
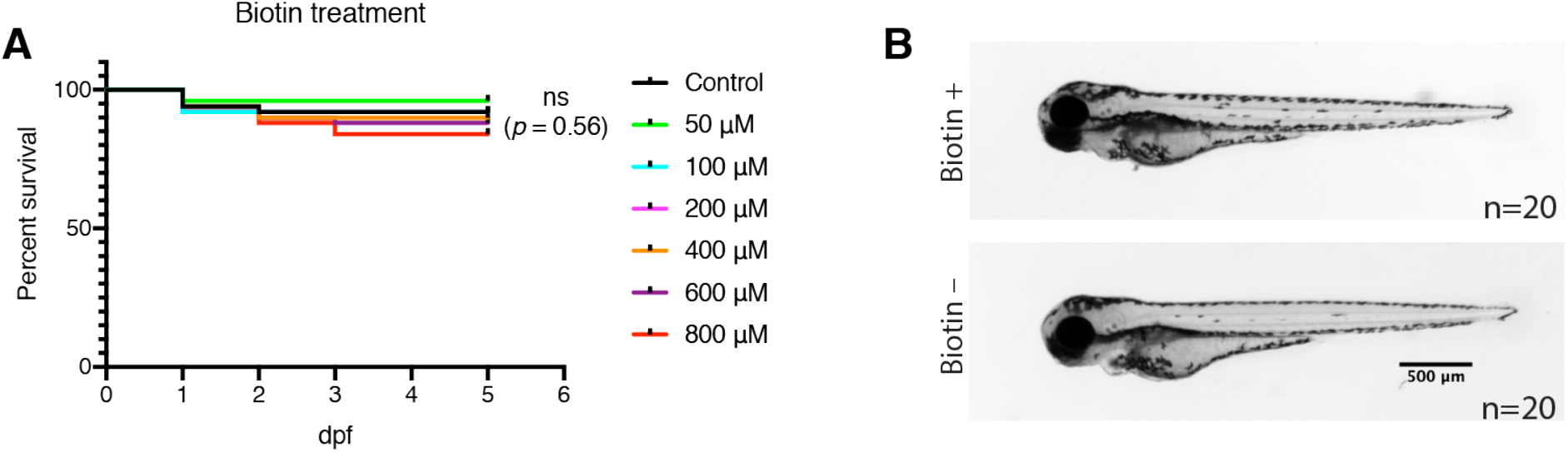
Determining biotin toxicity and tolerance in zebrafish embryos. (**A**) The fish embryos were incubated with biotin solution at indicated concentration from 1 to 5 dpf. Fifty WT embryos were used for each treatment group. Dead embryos were removed and counted each day. The statistics were performed using Log-rank (Mantel-Cox) test; ns denotes not significant. (**B**) representative images showing the fish embryo morphology at 3 dpf with or without biotin (600 μM) incubation.

**Figure S3.**
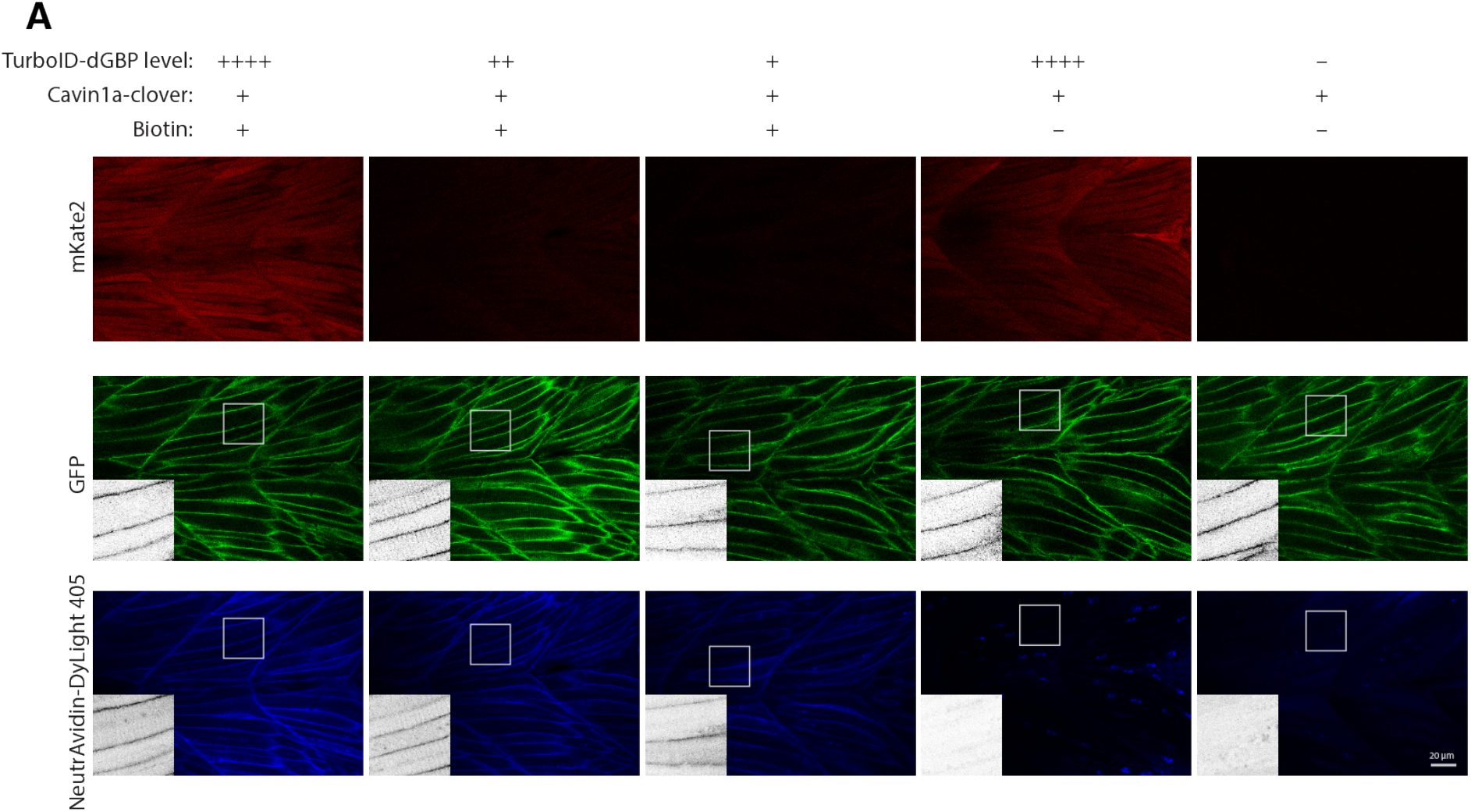
The specificity of TurboID-dGBP biotin labelling is independent of its expression level in zebrafish embryos. (**A**) Representative images showing biotinylated proteins in zebrafish embryos from outcrossing Cavin1a-Clover with different TurboID-dGBP lines. The number of “+” denotes the expression level of TurboID as reflected by mKate2 fluorescent reporter. The scale bar represents 20 μm.

**Figure S4.**
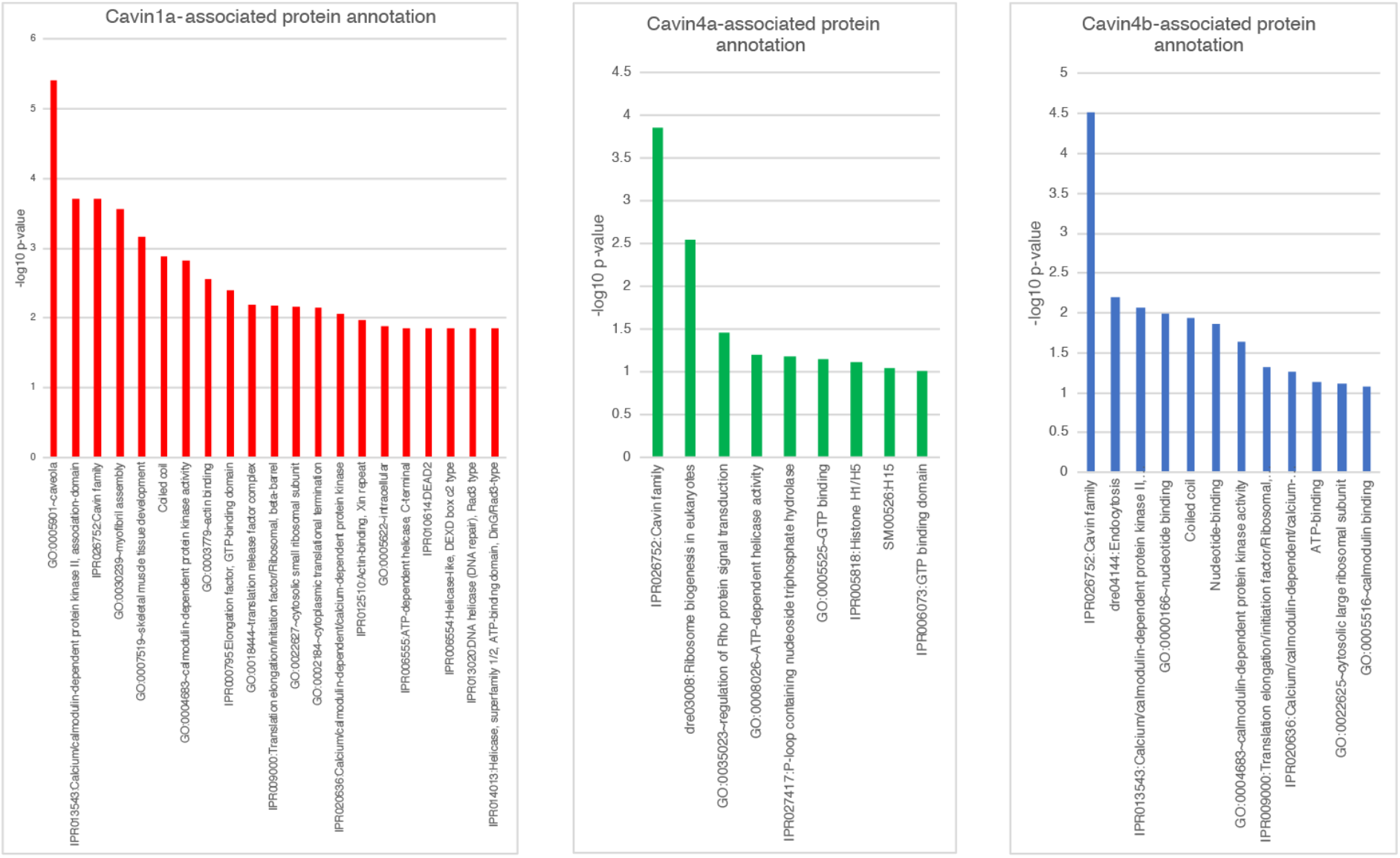
Gene ortholog annotation of cavin proteomes. The annotation was performed on DAVID Bioinformatics Resources 6.8. For proteins without existing information, their human or mouse orthologs were used. Y-axis shows −log10 adjust p-values from Benjamini-Hochberg procedure calculated by DAVID Bioinformatics Resources 6.8. Only proteins pass the cut-off (ES>0.9) were analysed.

